# Deep interpretable learning of sample representations for characterizing disease states in single-cell transcriptomics

**DOI:** 10.64898/2026.07.21.738207

**Authors:** Manoj M Wagle, Yongheng Wang, Soham Samanta, Zunpeng Liu, Ellis Patrick, Pengyi Yang, Manolis Kellis

## Abstract

Single-cell transcriptomics technology offers unprecedented insights into molecular heterogeneity. However, capturing sample-level representations that reflect both systemic and cellular states remains challenging, especially when disease annotations are mostly available as coarse sample-level labels. Here, we introduce Phenoverse, an interpretable deep learning framework that learns sample-level disease state representations through cell type-aware residual encoding, prototype learning, and Perceiver-based aggregation. Applied to independent single-cell transcriptomic cohorts of COVID-19, Alzheimer’s disease, and systemic lupus erythematosus, totaling over 5 million cells, we demonstrate that learned sample representations enable disease state prediction and encode a continuous spectrum of disease severity on unseen data that correlate with multiple clinical and pathological measures, despite being trained solely on binary phenotype labels. Further, we demonstrate that trajectory-derived genes reveal cross-cohort molecular programs and show consistently higher reproducibility than traditional case-control comparisons. Finally, prototype learning provides intrinsic model interpretability and enables the characterization of cell type-specific disease states. Taken together, Phenoverse offers an interpretable disease-phenotyping approach to dissecting sample heterogeneity, and our results highlight its utility in translating complex single-cell transcriptomic data into patient-level biological insights.

## Introduction

Over the past two decades, transcriptomic profiling has been widely used to interrogate the molecular state of patient samples and uncover gene programs underlying complex diseases. Bulk transcriptomics technologies enable us to capture such variation, but collapse the underlying cellular heterogeneity of samples into a single aggregate measurement. The establishment of single-cell transcriptomics addresses this limitation by recovering individual cell states across thousands of genes, providing a high-resolution view of phenotypic diversity (1–5). However, disease phenotypes are primarily annotated at the sample level and are often available only as coarse labels, such as disease versus control. Thus, translating high-resolution single-cell transcriptomic measurements into sample-level disease representations that capture biologically meaningful variation, including continuous disease severity and cellular disease states, remains an open problem.

Traditionally, sample-level analysis of single-cell data computes per-cell type average expression profiles or examines compositional changes in cell type abundance across samples (6). This condenses the data to a form equivalent to bulk measurements and discards the rich cellular heterogeneity of single-cell data. Furthermore, compositional analyses capture shifts in cell type abundance but do not account for transcriptional variation occurring within cell populations. Deep learning, a subfield of machine learning, is increasingly used in single-cell analyses due to its ability to handle high-dimensional and heterogeneous single-cell data (7–9). scVI introduced a probabilistic variational autoencoder (VAE) approach to model gene expression data into low-dimensional cell representations and has been shown to effectively handle batch effects, clustering, and other downstream tasks (10). More recently, foundation models pretrained on millions of cells have further extended this approach to provide cellular representations that generalize across different tissue types and conditions (11–14). Nevertheless, these methods do not account for sample-level context, and the inherent black-box nature of deep learning architectures offers limited insights into the cellular processes that drive their predictions (15–17).

Beyond the aforementioned challenges, the available disease annotations introduce further complexities in learning from single-cell data. Typically, such labels are predominantly binary annotations (e.g., disease vs control), and obtaining continuous clinical and pathological measures remains difficult. Case-control comparisons implicitly assume that the disease groups are homogeneous and can potentially hide important biological patterns, including disease severity gradients, molecularly distinct sample subgroups, and cell type-specific disease programs. Recent approaches have proposed learning multicellular sample-level representations from single-cell data. For example, PaSCient employs a DeepSet-based architecture and attention-based pooling to derive sample-level representations from individual expression profiles (18). mcBERT applies a transformer architecture with self-supervised pre-training followed by contrastive fine-tuning to generate sample representations (19). However, current methods are still limited in how well they capture the full spectrum of disease heterogeneity.

Here, we propose Phenoverse, an interpretable deep learning framework for characterizing sample-level disease states from single-cell transcriptomic data. Specifically, we adopt a three-component architecture, where (i) a cell type-aware residual neural network encoder models the gene expression data of individual cells, (ii) a prototype learning module decomposes cellular heterogeneity within each cell type into a set of learnable cellular states or prototypes, which are then concatenated with other cell type-level features to construct cell type tokens, and (iii) a Perceiver-inspired (20) cross attention module aggregates the cell type tokens into a sample-level disease representation (Fig. 1a). By learning only from binary phenotype labels, we show that across multiple diseases, Phenoverse representations enable disease state prediction, inference of clinically and pathologically aligned severity, and identification of biologically coherent disease subgroups. We demonstrate that the sample representations provide emergent interpretability through disease trajectories and facilitate the identification of reproducible cross-cohort molecular programs. In addition, prototype occupancies provide intrinsic concept-level interpretability and highlight disease-associated states within each cell type (Fig. 1b). Taken together, our results establish the utility of Phenoverse for translating high-resolution single-cell transcriptomic disease cohorts into sample-level disease continua and cell type-specific molecular programs.

**Fig. 1.**
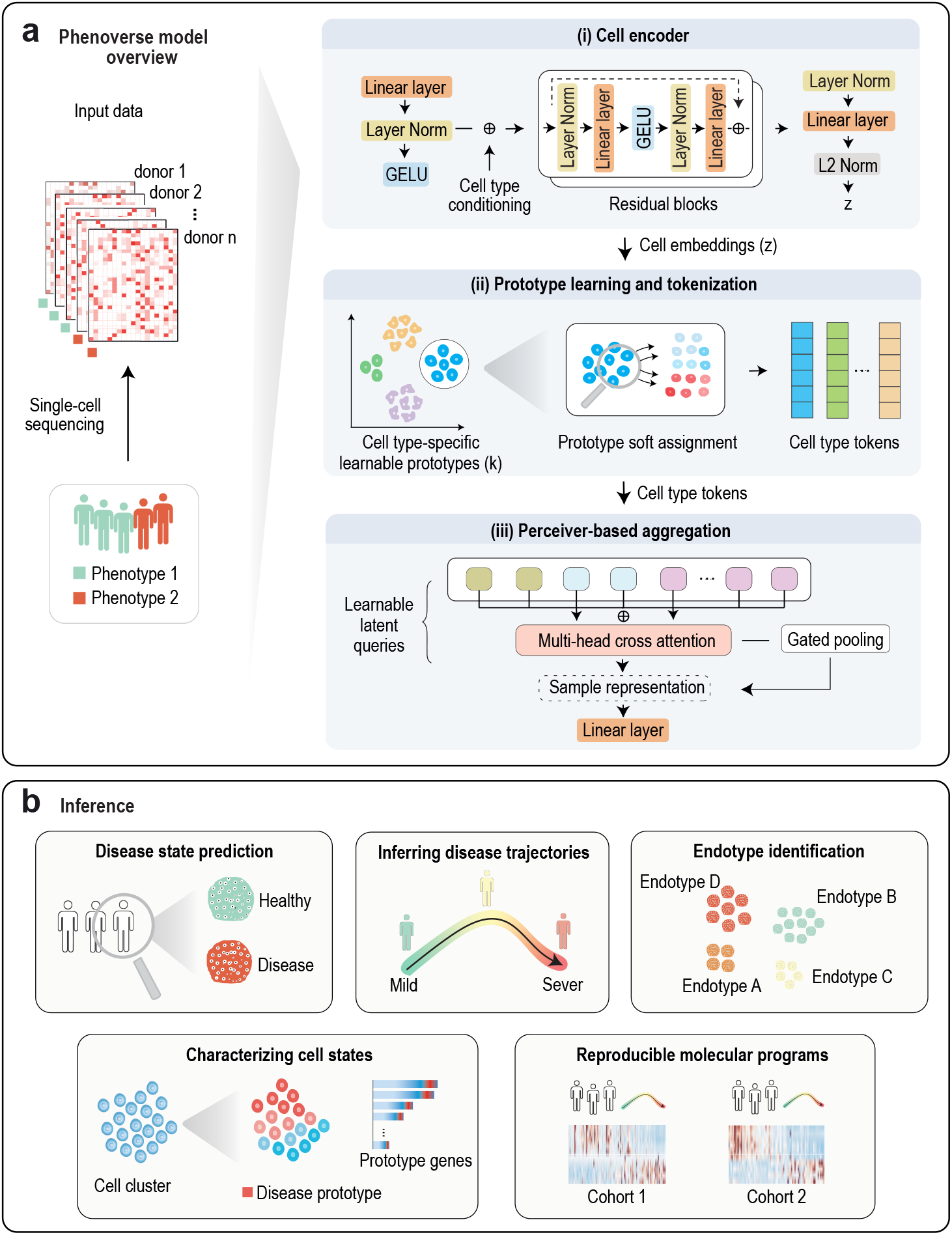
Overview of the Phenoverse framework for learning interpretable sample representations from single-cell transcriptomic data. (a) Graphical illustration of the Phenoverse model architecture with three jointly optimized components - a cell type-aware residual neural network encoder that generates *ℓ*_2_ normalized cell embeddings, a prototype learning and tokenization module that softly assigns cells to learnable cell type-specific prototypes, which are then concatenated with cell type-level features to construct cell type tokens, and a Perceiver-based aggregator that applies multi-head cross-attention between learnable latent queries and cell type tokens to produce the sample representation. (b) Schematic of downstream applications of Phenoverse, including disease state prediction for unlabeled samples, inferring continuous disease trajectories, identifying disease subgroups, characterizing cell states through prototype gene programs, and discovering reproducible molecular programs across independent cohorts.

## Results

### Overview of model

We developed Phenoverse as an interpretable disease-phenotyping model that learns from individual cell expression profiles to produce sample-level disease state representations and characterize heterogeneity from single-cell transcriptomic cohorts. The model processes each sample through three components that are jointly optimized end-to-end (Fig. 1a). First, we map the high-dimensional gene expression profile of individual cells into a low-dimensional embedding using a residual neural network. Cell type identity is incorporated by adding a learnable cell type embedding. The prototype pooling and tokenization module then processes cell embeddings for each cell type by employing gated attention to weight the contribution of each cell. An attention-weighted mean and standard deviation are then derived for each cell type. Following this, each cell is softly assigned to learnable prototype vectors through temperature-scaled cosine similarity. Next, the mean state, standard deviation, prototype occupancy, and cell type proportion are projected through a multi-layer perceptron, and a cell type token is obtained for each cell type. Finally, a Perceiver-based aggregator uses learnable latent queries to attend to these tokens via cross-attention. The resulting outputs are pooled into a sample representation and passed through a linear classification head (Methods).

### Predicting disease state for unlabeled samples

We first evaluated the quality of the sample representations by examining whether they are discriminative for disease state prediction. We applied Phenoverse to a large-scale single-cell transcriptomic atlas from COVID-19, comprising approximately 1.4 million cells across approximately 182 samples (21). Using a representative held-out split of samples, we found that annotated cell types showed clear separation in the cell latent space (Fig. 2a). Similarly, the two classes ‘COVID-19’ and ‘Normal’ exhibited a clear separation in the sample representation space (Fig. 2b). Next, we quantified the sample disease prediction performance using a five-time repeated subsampling validation against seven baseline methods, including sample representation methods - PaSCient (18) and mcBERT (19), representations derived from single-cell foundation models - scGPT (11) and tGPT (13), a deep generative model - scVI (10), simple multilayer perceptron baseline (Simple MLP), and principal component analysis (PCA) (Methods).

**Fig. 2.**
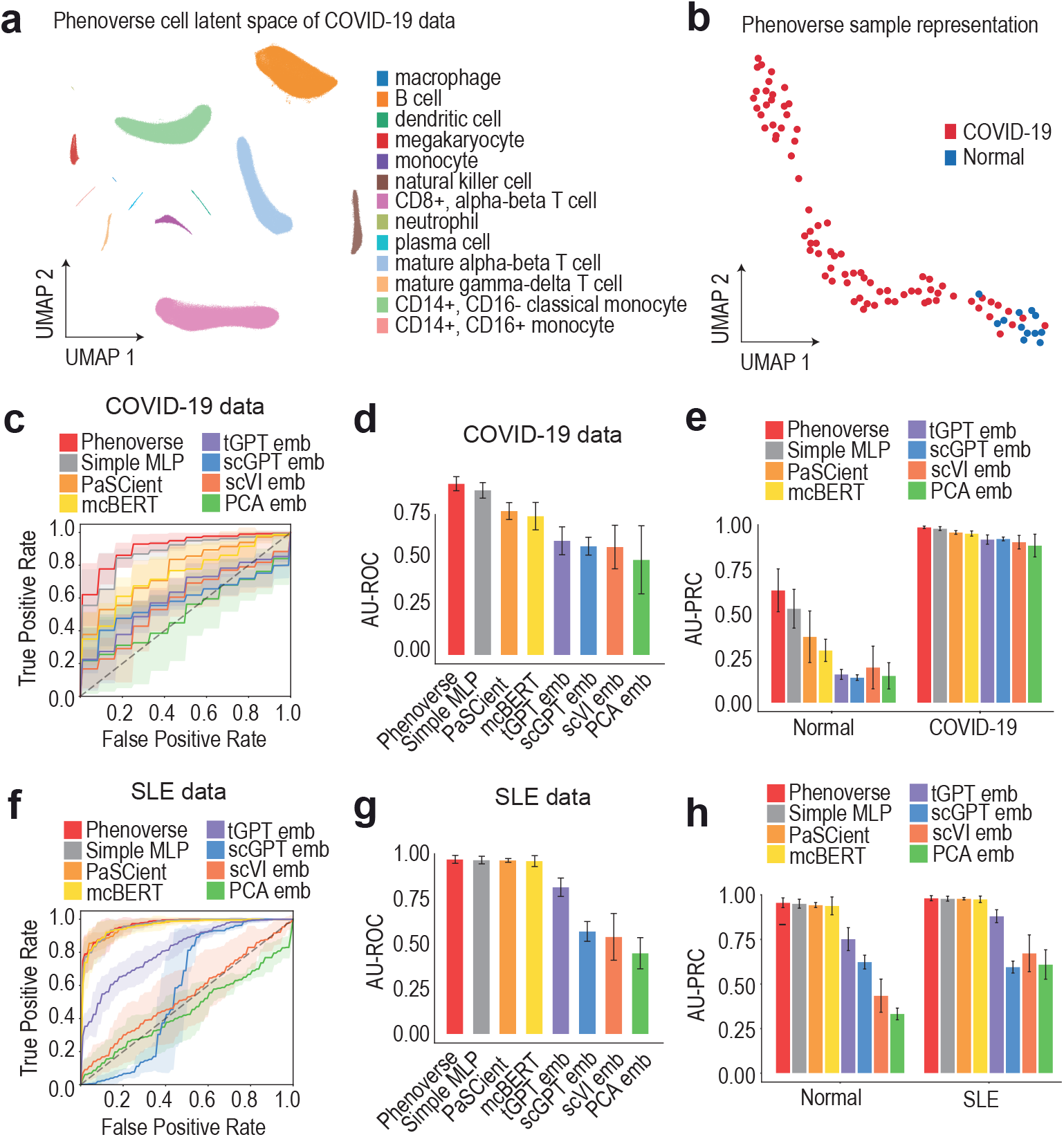
Evaluation of sample representations for disease state prediction in unlabeled data. (a) Uniform manifold approximation and projection (UMAP) of the Phenoverse cell embeddings from a representative held-out test set of the COVID-19 dataset colored by cell type. (b) UMAP of Phenoverse sample representations from a representative held-out test set of the COVID-19 dataset, colored by disease state. (c) Mean receiver operating characteristic curve (ROC) with *±*1 standard deviation shaded bands illustrating five-time repeated subsampling validation performance for disease state prediction in the COVID-19 dataset. (d-e) Bar plots showing sample disease prediction performance using the area under the receiver operating characteristic curve (AU-ROC) (d) and per-class area under the precision-recall curve (AU-PRC) (e) metrics in the COVID-19 dataset. Error bars represent the standard deviation across five repeated subsampling runs. ‘_emb’ denotes sample representations derived from cell embeddings. (f) Mean receiver operating characteristic curve (ROC) with *±*1 standard deviation shaded bands illustrating five-time repeated subsampling validation performance for disease state prediction in the systemic lupus erythematosus (SLE) dataset. (g-h) Bar plots showing sample disease prediction performance using the area under the receiver operating characteristic curve (AU-ROC) (g) and per-class area under the precision-recall curve (AU-PRC) (h) metrics in the systemic lupus erythematosus (SLE) dataset. Error bars represent the standard deviation across five repeated subsampling runs. ‘_emb’ denotes sample representations derived from cell embeddings.

Phenoverse achieved an overall area under the receiver operating characteristic curve (AU-ROC) of 0.914 *±* 0.038, outperforming baseline methods (Fig. 2c-d). The Simple MLP baseline performed similarly to our method and ranked second. To account for class imbalance, we further evaluated performance using the area under the precision-recall curve (AU-PRC). All methods performed mostly similarly in the ‘COVID-19’ class, while Phenoverse achieved substantially higher performance compared to other methods in the ‘Normal’ class (Fig. 2e, Supplementary Fig. 1a). We next evaluated Phenoverse on another single-cell transcriptomic cohort from systemic lupus erythematosus (SLE), comprising approximately 1.2 million cells across approximately 261 samples (22). Using five-time repeated subsampling validation, Phenoverse achieved an overall AU-ROC of 0.969 *±* 0.022 with PaSCient, mcBERT, and Simple MLP attaining a comparable performance (Fig. 2f-g). Similarly, for AU-PRC, Phenoverse, PaSCient, mcBERT, and Simple MLP achieved consistently high performance in both classes (Fig. 2h, Supplementary Fig. 1b). In summary, these results demonstrate that Phenoverse sample representations retain discriminative information and accurately predict disease states across independent datasets.

### Inferring clinically and pathologically aligned disease severity with Phenoverse

We next examined whether the sample representations can recapitulate the continuous spectrum of disease severity beyond the binary labels. COVID-19 is an infectious disease exhibiting marked clinical heterogeneity, ranging from mild to moderate illness to life-threatening disease. Thus, accurately staging each patient along this spectrum is important for clinical outcome prediction and treatment decisions (23–25). We used the COVID-19 single-cell transcriptomic dataset, which consists of three clinical severity categories - control, mild/moderate, and severe/critical (21). We used the five-time repeated subsampling validation to obtain different variants of held-out data. The disease trajectories from the sample representations of each method were inferred using diffusion pseudotime anchored at the control sample (Methods).

We found that Phenoverse sample representations outperformed baseline methods and recovered the clinical severity with an overall Spearman correlation of 0.48 (Fig. 3a-d, Supplementary Fig. 2a-b, Methods). Simple MLP baseline ranked second, while representations derived from scGPT, PCA, and scVI outperformed both sample representation methods - mcBERT and PaSCient. To further characterize the inferred trajectory, we fit an ordinal logistic regression model to the Phenoverse pseudotime values (Methods). Phenoverse showed an odds ratio of 28.11 (*p* = 4.6 *×* 10−25), suggesting a strong association between the inferred trajectory and disease severity. We found that the probability of being classified as control decreased monotonically along the trajectory, the probability of mild/moderate peaked at intermediate pseudotime values, and the probability of severe/critical increased monotonically (Fig. 3e, Supplementary Fig. 2c). Next, we evaluated Phenoverse using our single-cell transcriptomic data on Alzheimer’s disease of the aged prefrontal cortex (Religious Orders Study and Rush Memory and Aging Project - ROSMAP cohort), comprising approximately 1.4 million cells across approximately 294 samples (26). Alzheimer’s disease (AD) is a neurodegenerative disorder characterized by the accumulation of abnormal proteins (amyloid plaques and neurofibrillary tangles), resulting in cognitive decline and widespread neuronal cell death (27–30). Using the same five-time repeated subsampling validation approach, we inferred disease trajectories from the sample representations anchored at samples with the lowest neuropathological burden (Methods).

**Fig. 3.**
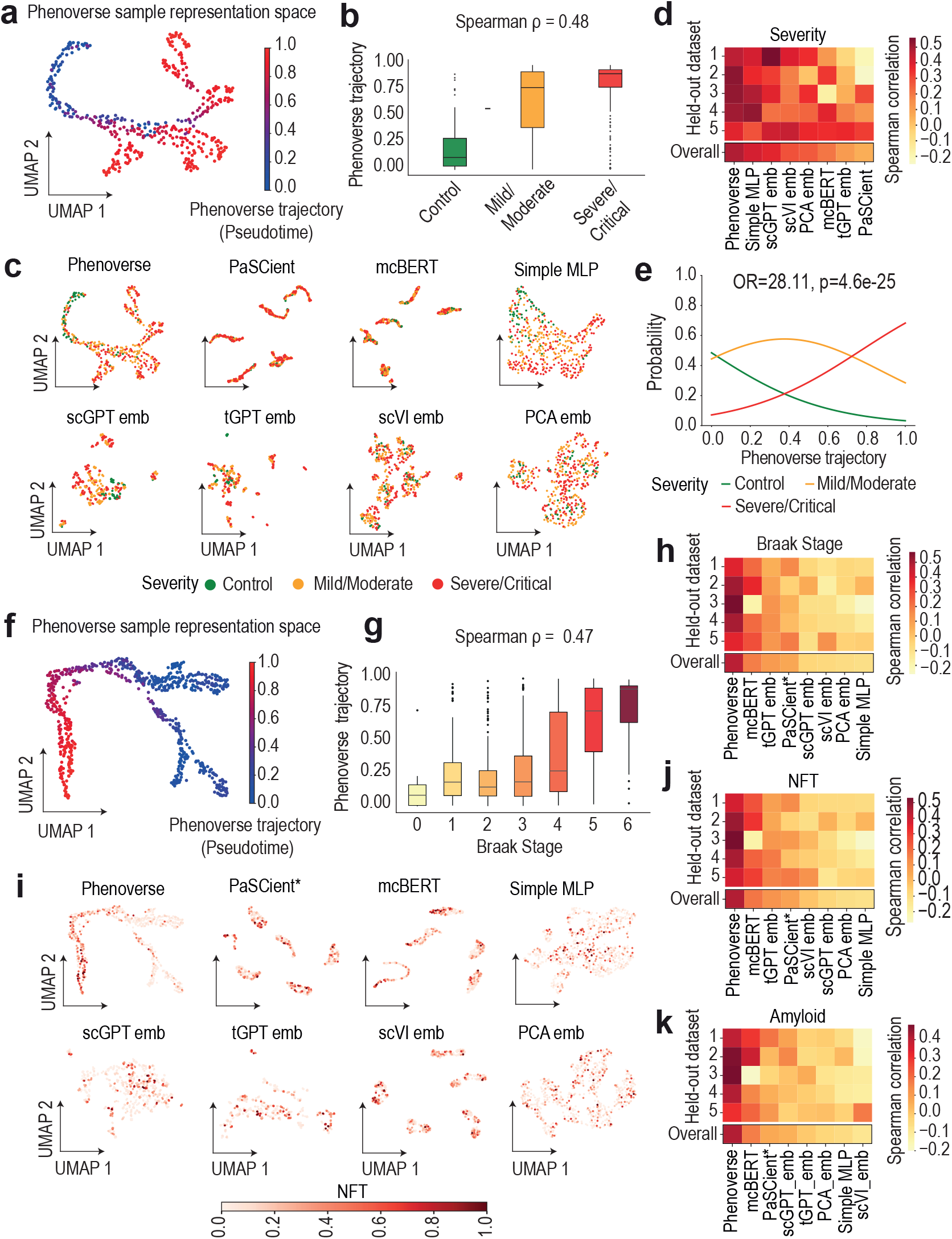
Evaluation of inferred disease trajectories. (a) UMAP of Phenoverse sample representations from held-out test samples in the COVID-19 dataset, colored by inferred trajectory (pseudotime). (b) Box plot showing the distribution of Phenoverse inferred trajectories across clinical severity categories (control, mild/moderate, and severe/critical) in the COVID-19 dataset. Box plots show the median (center line), interquartile range (IQR) box bounds: 25th–75th percentile, 1.5*×*IQR (whiskers), and points outside 1.5*×*IQR as outliers. (c) UMAPs of sample representations of different methods from held-out test samples in the COVID-19 dataset, colored by clinical severity. Heatmap showing the Spearman correlation between inferred trajectory and clinical severity across five repeated subsampling validation splits in the COVID-19 dataset. Line plot showing ordinal logistic regression predicted probabilities for each clinical severity category along the Phenoverse inferred trajectory in the COVID-19 dataset. Odds ratios (OR) and associated p-values are indicated in the panel title. (f) UMAP of Phenoverse sample representations from held-out test samples in the ROSMAP Alzheimer’s disease cohort, colored by inferred trajectory (pseudotime). (g) Box plot showing the distribution of Phenoverse inferred trajectories across Braak stages (0–6) in the ROSMAP data. Box plots show the median (center line), interquartile range (IQR) box bounds: 25th–75th percentile, 1.5*×*IQR (whiskers), and points outside 1.5*×*IQR as outliers. (h) Heatmap showing the Spearman correlation between inferred trajectories and Braak stage across five repeated subsampling validation splits for different methods in the Alzheimer’s disease dataset. * indicates that pseudotime values were undefined for a subset of held-out samples. (i) UMAPs of sample representations from different methods for held-out test samples in the ROSMAP cohort, colored by neurofibrillary tangles (NFT) burden. * indicates that pseudotime values were undefined for a subset of held-out samples. (j-k) Heatmaps showing the Spearman correlation between inferred trajectory and NFT burden (j) and amyloid burden (k) across five repeated subsampling validation splits in the ROSMAP data. * indicates that pseudotime values were undefined for a subset of held-out samples.

We found that the sample representation space of Phenoverse revealed a continuous trajectory that aligned with increasing pathological burden, and showed a Spearman correlation of 0.47 with the Braak stage (Fig. 3f-g). In contrast, most baseline methods failed to recover meaningful correlations with Braak staging, with the second-best performing method, mcBERT, achieving a Spearman correlation of 0.12 (Fig. 3h, Supplementary Fig. 3a-b). We further evaluated the inferred trajectories of different methods against additional pathological measures, including neurofibrillary tangles (NFT) and amyloid burden. Again, Phenoverse achieved the highest correlation (NFT burden: Spearman *ρ* = 0.46, Pearson *ρ* = 0.52; Amyloid burden: Spearman *ρ* = 0.40, Pearson *ρ* = 0.44), outperforming other methods (Fig. 3i-k, Supplementary Fig. 4a-b). Simple MLP baseline, despite being ranked second in the COVID-19 cohort, performed worst among baseline methods across two pathological measures (Braak stage and NFT) in Alzheimer’s disease. Thus, by learning only from binary phenotype labels, Phenoverse sample representations encode a continuous spectrum of disease severity that aligns well with multiple clinical and pathological measures of disease progression.

### Phenoverse trajectories reveal reproducible cross-cohort molecular programs in Alzheimer’s disease

Alzheimer’s disease presents as a progressive biological continuum characterized by varying degrees of neuropathological accumulation. Standard case-control comparisons often collapse this heterogeneity and could miss molecular programs that shift along the disease trajectory. Using the ROSMAP data, we trained Phenoverse and applied it to held-out samples comprising approximately 700,000 cells across 147 samples and seven cell types (26). We found that the cell latent space showed a clear separation of cell types, and cells from AD and nonAD groups were segregated within each cell type (Fig. 4a). Similarly, the representation space of samples appeared separated by disease groups (Fig. 4b). The disease trajectory was then inferred from the sample representations, which showed a Spearman correlation of 0.37 with amyloid burden (Fig. 4c, Supplementary Fig. 5a). Next, we used the same trained model and applied it to another independent single-cell expression cohort from the Seattle Alzheimer’s Disease Brain Cell Atlas (SEA-AD). This dataset consists of approximately 1.4 million cells across 83 samples from the dorsolateral prefrontal cortex (Fig. 4d-e, Supplementary Fig. 5a) (31). The disease trajectory was then inferred and correlated with the CERAD (Consortium to Establish a Registry for Alzheimer’s Disease) score, a pathological measure that quantifies the density of neuritic amyloid plaques. We found that the inferred trajectory showed a Spearman correlation of 0.53 with the CERAD score, demonstrating the generalizability of Phenoverse in capturing disease progression (Fig. 4f).

**Fig. 4.**
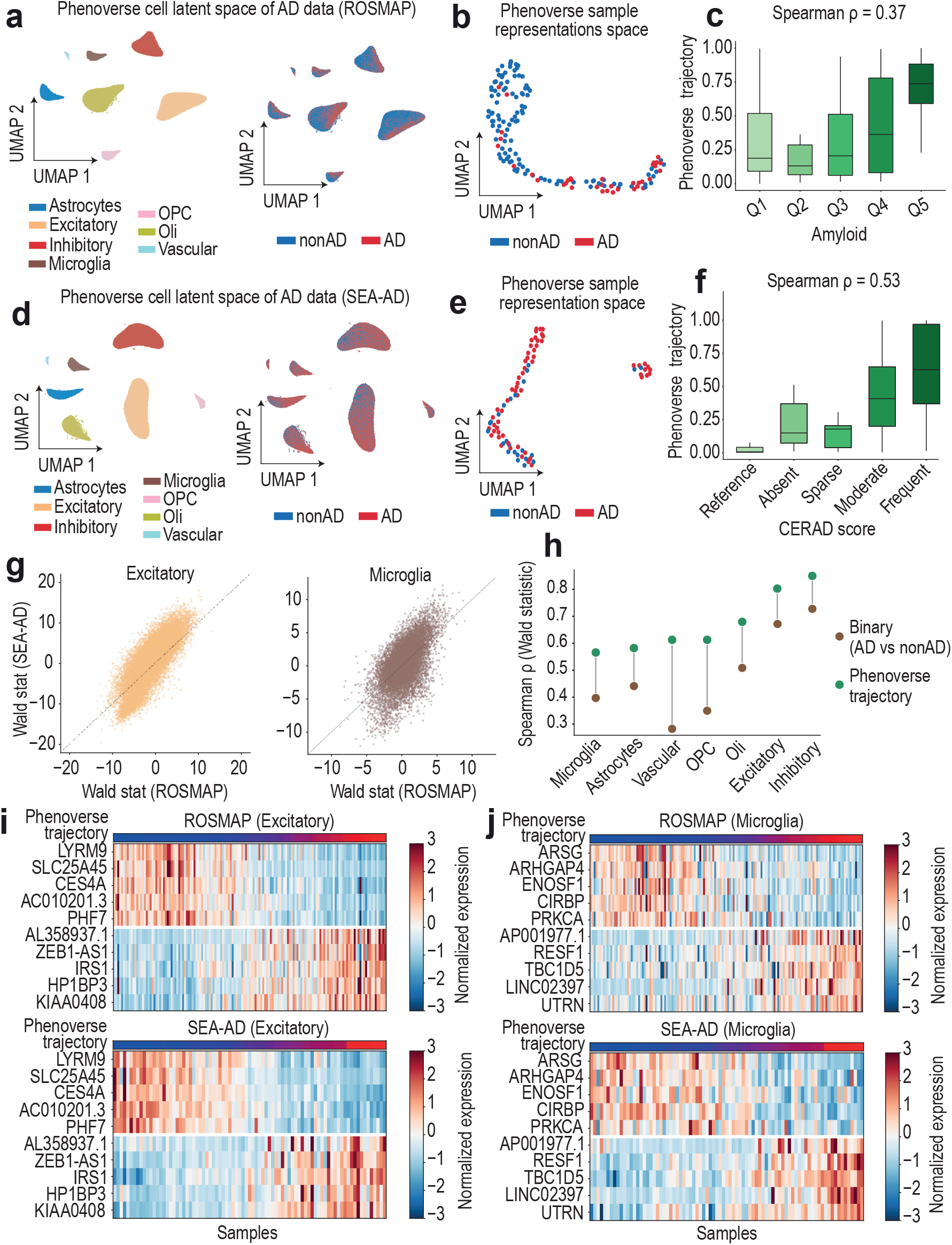
Identifying cross-cohort molecular programs in Alzheimer’s disease with Phenoverse. (a) UMAPs of Phenoverse cell embeddings from held-out samples of the ROSMAP cohort, colored by cell type (left) and disease group (right). (b) UMAP of Phenoverse sample representations from held-out samples of ROSMAP cohort, colored by disease group. (c) Box plot showing the distribution of Phenoverse inferred trajectory values across amyloid burden quantiles (Q1–Q5) in the ROSMAP cohort. Box plots show the median (center line), interquartile range (IQR) box bounds: 25th–75th percentile, 1.5*×*IQR (whiskers), and points outside 1.5*×*IQR as outliers. (d) UMAPs of Phenoverse cell embeddings from the SEA-AD cohort, colored by cell type (left) and disease group (right). (e) UMAP of Phenoverse sample representations from held-out samples of SEA-AD cohort, colored by disease group. (f) Box plot showing the distribution of Phenoverse inferred trajectory values across CERAD score categories in the SEA-AD cohort. Box plots show the median (center line), interquartile range (IQR) box bounds: 25th–75th percentile, 1.5*×*IQR (whiskers), and points outside 1.5*×*IQR as outliers. (g) Scatter plots showing the cross-cohort reproducibility of gene-level Wald statistics from trajectory-based differential expression analysis in excitatory neurons (left) and microglia (right). (h) Dumbbell plot comparing the cross-cohort reproducibility of gene-level differential expression Wald statistics between ROSMAP and SEA-AD cohorts across cell types. (i-j) Heatmaps illustrating expression patterns of the top upregulated and downregulated trajectory-associated genes along the Phenoverse trajectory in Excitatory neurons (i) and Microglia (j). Genes were selected based on Wald statistics from the ROSMAP cohort. Expression values are shown as row-normalized Z-scores. Rows correspond to genes, and columns correspond to samples ordered by increasing pseudotime.

Next, we independently characterized the transcriptomic changes underlying the inferred trajectories in both cohorts by performing differential expression analysis with pseudotime as a continuous covariate (32, 33). Additionally, for each cell type, binary pseudobulk differential expression analysis was performed using the available disease group labels in each cohort (Methods). We found that trajectory-derived genes showed consistently higher cross-cohort reproducibility across all cell types compared to binary case-control comparison (Fig. 4g-h, Supplementary Fig. 5b). Further, we used top upregulated and downregulated trajectory-associated genes from the ROSMAP cohort and examined their expression patterns along the inferred trajectory in both cohorts. The genes showed clear progressive changes in expression along the trajectory in ROSMAP samples, and a similar expression pattern was observed in the SEA-AD cohort (Fig. 4i-j, Supplementary Fig. 6a). Finally, the pathway activity scores were correlated with the Phenoverse inferred trajectory to characterize shared molecular programs in both cohorts (Supplementary Fig. 7a, Methods) (34–37). We found that inflammatory response, G2-M checkpoint, and TNF-alpha signaling via NF-kB pathways were enriched in neuronal populations (excitatory and inhibitory) and oligodendrocytes. This is consistent with the NF-kB-mediated neuronal stress, and aberrant cell cycle re-entry in post-mitotic neurons (38, 39). Further, epithelial mesenchymal transition was progressively enriched in excitatory neurons, while oxidative phosphorylation and interferon alpha response pathways were depleted in inhibitory neurons (40–42). We also found that heme metabolism was enriched across most cell types, potentially reflecting the widespread iron dysregulation and impaired heme turnover associated with mitochondrial dysfunction (43, 44). In astrocytes, cholesterol homeostasis was depleted, and PI3K/AKT/mTOR signaling was enriched, whereas microglia showed enrichment of androgen response programs (45–47). Finally, in vascular cells, the reactive oxygen species and p53 pathways were progressively enriched, aligning with blood-brain barrier dysfunction due to oxidative stress and endothelial DNA damage (48, 49). These results show that Phenoverse-inferred trajectories reveal meaningful cell type-resolved molecular programs of disease progression in Alzheimer’s disease.

### Phenoverse learns biologically coherent disease states in Systemic Lupus Erythematosus

Systemic lupus erythematosus (SLE) or lupus is a chronic autoimmune disease characterized by dysregulation of interferon signaling in circulating immune cell compartments, and the underlying molecular drivers of this heterogeneity remain poorly understood (50–52). Here, using the single-cell transcriptomic PBMC dataset, we investigated whether Phenoverse can identify cell type-specific disease states by learning from binary sample phenotype labels (22). We trained Phenoverse and applied it to held-out samples comprising approximately 600,000 cells across 131 samples spanning nine cell types (Fig. 5a). We found that the representation space exhibited a clear separation of samples between SLE and healthy groups (Fig. 5b). Next, to assess whether the model captures disease-relevant variation, we clustered the sample representations of held-out samples and identified seven distinct groups. For each sample, we then computed an IFN score using interferon-stimulated gene expression markers. SLE-enriched clusters showed elevated IFN activity relative to healthy-enriched clusters, and IFN scores increased progressively from healthy to disease-enriched clusters. This monotonic increase in IFN activity confirms that Phenoverse sample representations organize along a continuous disease axis and stratify into biologically coherent endotypes (Fig. 5c–d).

**Fig. 5.**
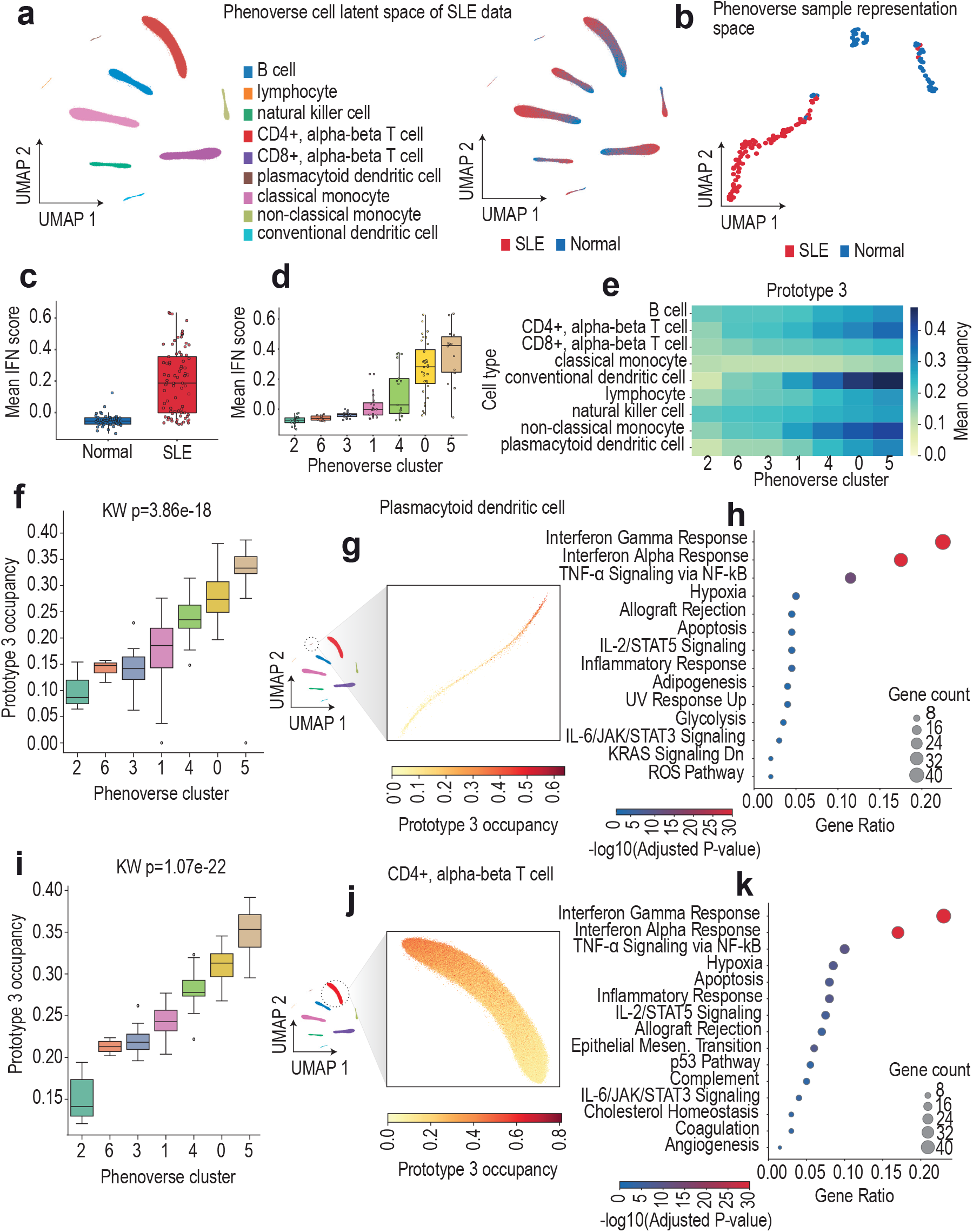
Phenoverse identifies cell type-specific disease states in Systemic Lupus Erythematosus. (a) UMAPs of Phenoverse cell embeddings from held-out samples, colored by cell type (left) and disease group (right). (b) UMAP of Phenoverse sample representations, colored by disease group. (c) Boxplots showing distribution of per-donor IFN scores stratified by disease group. Box plots show the median (center line), interquartile range (IQR) box bounds: 25th–75th percentile, 1.5*×*IQR (whiskers), and points outside 1.5*×*IQR as outliers. (d) Boxplots illustrating mean per-sample IFN scores across the seven Phenoverse clusters ordered by increasing IFN score. (e) Heatmap illustrating prototype 3 occupancy profiles across sample clusters for different cell types, ordered by increasing IFN score. (f) Box plots showing the distribution of prototype 3 occupancy across Phenoverse sample clusters in plasmacytoid dendritic cells, ordered by increasing IFN score. Box plots show the median (center line), interquartile range (IQR) box bounds: 25th–75th percentile, 1.5*×*IQR (whiskers), and points outside 1.5*×*IQR as outliers. (g) UMAP highlighting prototype 3 occupancy in plasmacytoid dendritic cells. (h) Dot plot illustrating enrichment of top prototype 3 gene programs in plasmacytoid dendritic cells. (i) Box plots showing the distribution of prototype 3 occupancy across Phenoverse sample clusters in CD4-positive T cells, ordered by increasing IFN score. Box plots show the median (center line), interquartile range (IQR) box bounds: 25th–75th percentile, 1.5*×*IQR (whiskers), and points outside 1.5*×*IQR as outliers. (j) UMAP highlighting prototype 3 occupancy in CD4+ alpha-beta T cells. (k) Dot plot illustrating enrichment of top prototype 3 gene programs in CD4+ alpha-beta T cells.

We next asked whether Phenoverse can identify cell type-specific disease states by interpreting the prototype occupancies across different clusters (Fig. 5e). Across all nine cell types, there was a consistent increase in prototype 3 (P3) occupancy from healthy-to SLE-enriched clusters (Fig. 5e). We characterized the gene programs underlying these shifts by computing correlations between per-cell gene expression and per-cell prototype occupancy. Enrichment analysis was then performed on the top 200 positively correlated genes against hallmark gene sets from the Molecular Signatures Database (MSigDB) (Methods). We first examined plasmacytoid dendritic cells (pDC), the dominant type I interferon-secreting cell type in systemic lupus erythematosus (Fig. 5g-h) (53, 54). The prototype 3 gene program was found to be strongly enriched for interferon alpha response, interferon gamma response, and TNF-alpha signaling via NF-kB, consistent with the known activated state of plasmacytoid dendritic cells in SLE (Fig. 5h) (55, 56). We next validated these findings in another representative cell type, CD4+ alpha-beta T cells, which are known to regulate IFN-stimulated genes and adopt pro-inflammatory effector states in SLE (Fig. 5i-j) (57, 58). The prototype 3 gene programs in CD4+ alpha-beta T cells were again enriched for interferon response, TNF-alpha signaling via NF-kB, and inflammatory response pathways. Similarly, across the remaining seven cell types, the P3 gene programs consistently reflected an interferon-driven enrichment. These results show that, through learned prototype-based interpretation, Phenoverse enables characterization of cell disease states and the molecular programs underlying systemic lupus erythematosus.

## Discussion

Here, we introduced Phenoverse as an interpretable deep learning framework for learning sample-level disease state representations from coarsely labeled single-cell transcriptomic data. The framework jointly trains a cell type-aware residual encoder, a prototype pooling and tokenization module, and a Perceiver-based cross-attention aggregator to generate a unified representation for each sample. By incorporating learnable prototypes into the model architecture, Phenoverse provides intrinsic interpretability and enables the delineation of cellular disease states.

Despite being trained only on binary phenotype labels, Phenoverse sample representations revealed a continuous disease axis that aligned with independent measures of disease severity. In COVID-19, the inferred trajectories recovered clinical progression from mild to severe disease. In Alzheimer’s disease, the trajectories correlated with multiple independent measures of neuropathological burden, including Braak stage, amyloid, and neurofibrillary tangles, outperforming baseline approaches. Further, trajectory-derived differential expression rankings revealed cross-cohort molecular programs and showed consistently higher reproducibility compared to binary case-control comparison across all cell types. In systemic lupus erythematosus, the sample representation space stratified along a gradient of interferon activity, forming biologically coherent endotypes. Finally, gene programs characterized through disease-associated prototypes were predominantly enriched for interferon-driven pathology in SLE. These findings show that prototype-based learning provides meaningful insights into cell type-specific programs underlying disease heterogeneity.

In this paper, we applied Phenoverse across diverse disease contexts, however, we acknowledge that our framework is currently limited to learning from single-cell transcriptomic data. Future extensions of this work could therefore include incorporating multimodal single-cell measurements, including chromatin accessibility and surface proteins, as well as spatial omics. This would enable a multi-layered characterization of disease states by integrating regulatory, proteomic, and spatial dimensions of cellular variation. As omics technologies continue to advance, we anticipate that sample representation approaches will become central to translating high-dimensional single-cell measurements into valuable patient-level insights.

## Methods

### Model architecture

Phenoverse processes each sample as a set of *n* cells {**x**_1_, … , **x**_*n*_ , where **x**_*i*_ ∈ ℝ*G* denotes the gene expression profile of the cell *i* over genes *G*. The corresponding cell type label of the cell is denoted as *c*_*i*_ ∈ {1,…, *C}* . We start with a raw single-cell count matrix that is log-transformed and scaled:

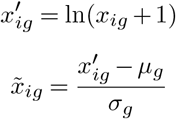

where *x*_*ig*_ is the raw count for cell *i* and gene *g, µ*_*g*_ and *σ*_*g*_ are the mean and standard deviation of gene *g* across all cells. To address sample-level class imbalance, we dynamically construct sample bags at each epoch by oversampling minority-class samples with replacement to match the majority-class sample count. Further, at each training step, we subsample 500-1000 cells per sample stratified by cell types. We then feed the processed data into the model that integrates three components trained end-to-end, including a cell type-aware residual encoder, a per-cell type prototype pooling and tokenization module, and a Perceiver-based cross-attention aggregator.

#### Cell encoder

First, we employ a cell type-aware residual neural network encoder to generate cell embeddings. For a given cell *i* with expression profile **x**_*i*_, the input is passed through a linear layer with weight matrix 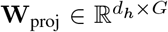 and bias 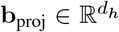. This is followed by layer normalization and GELU activation (59). We incorporate cell identity into the model by adding a learnable cell type embedding 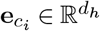:

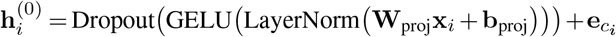

The representation is further refined through residual blocks (60), each comprising two pre-normalized linear transformations with GELU activation and dropout. The output is finally projected through a linear layer with weight matrix 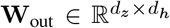 and bias 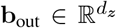. This is subsequently *ℓ*_2_ normalized to produce the cell embedding 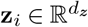:

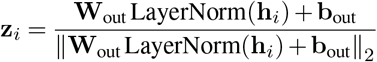

#### Prototype pooling and tokenization

For each cell type *c* ∈ {1,…, *C}* , let *Ƶ*_*c*_ = {**z**_*i*_ : *c*_*i*_ = *c}* denote the cell embeddings. We first compute a scalar score for each cell using a gated attention multi-layer perceptron MLP_gate_, comprising layer normalization followed by two linear transformations and GELU activation. The scores are then normalized via softmax over all cells of type *c* to obtain per-cell attention weights *α*_*i*_:

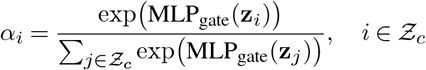

The attention-weighted mean state ***µ****c* and element-wise standard deviation ***σ***_*c*_ are then computed as:

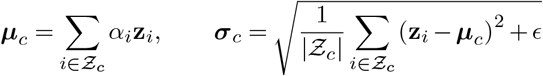

where *ϵ* is a constant. We assign each cell to a set of *K* learnable prototype vectors 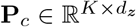, initialized using the Xavier uniform initialization (61). The soft assignment to a prototype is computed through temperature-scaled cosine similarity between the cell embeddings and prototypes:

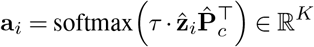

where *τ* is a learnable temperature parameter. The prototype occupancy **o**_*c*_ ∈ ℝ*K* , representing the mean assignment of cells across prototypes is given by:

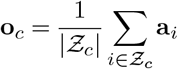

Finally, a cell type token 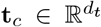 is obtained by concatenating the mean state, standard deviation, prototype occupancy, cell type proportion *π*_*c*_ = |*Ƶ*|_*c*_ */n*, and intra-cell type variability *η*_*c*_ = mean(***σ***_*c*_). This is then passed through a token multi-layer perceptron with a residual cell type embedding 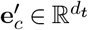.

#### Perceiver-based aggregator

The cell type tokens 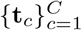 obtained from the previous step are aggregated into a unified sample representation using a Perceiver-inspired cross-attention architecture (20). We introduce a set of *M* learnable latent query vectors 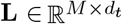, initialized using the Xavier uniform initialization. These queries then attend to the cell type tokens 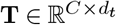 via cross-attention. We then pool the learned latents into a single sample representation 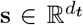. Specifically, a scalar weight *β*_*l*_ is assigned to each latent through a gating multi-layer perceptron and normalized with softmax:

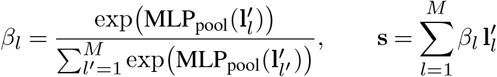

The sample representation **s** is then passed through a linear classification head with a cross-entropy phenotype classification loss. We hold out 20% of the training data as a validation subset, stratified by phenotype label. The validation data is used to monitor model training, and an early stopping mechanism is implemented based on the validation classification loss, with a patience of ten epochs. The classification loss is backpropagated and all model parameters are jointly updated, including the cell type Embeddings 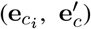, prototype vectors (**P**_*c*_), and latent queries (**L**), using AdamW optimization (62).

### Baseline methods

#### PaSCient

PaSCient was implemented following the authors’ official documentation (https://github.com/genentech/pascient). Raw count matrices were normalized to counts per 10000 and log1p-transformed. Class-balanced oversampling was applied at each training epoch using the OverSamplerPerAttribute function. The model was trained with the following parameters - ‘lr’ = 1e-4, ‘weight_decay’ = 1e-4, ‘max_epochs’ = 100, ‘batch_size’ = 16, ‘sample_size’ = 1500, and ‘patience’ = 10. Sample representations were obtained for held-out samples using the trained model.

#### mcBERT

mcBERT was implemented following the authors’ official documentation (https://github.com/COMSYS/mcBERT). A total of 1000 highly variable genes were selected from the training data using the Seurat v3 method. The raw count matrices were normalized to total counts. Pre-training was performed using the data2vec self-supervised objective with the following parameters - ‘embed_dim’ = 288, ‘num_hidden_layers’ = 12, ‘num_attention_heads’ = 12, ‘mlm_probability’ = 0.15, ‘batch_size’ = 8, ‘lr’ = 1e-5, ‘weight_decay’ = 0.01, and ‘num_epochs’ = 10. The pre-trained encoder was then fine-tuned using a supervised contrastive loss with the following parameters: ‘temperature’ = 0.1, ‘n_cells’ = 1023, ‘lr’ = 1e-5, ‘weight_decay’ = 0.01, ‘num_epochs’ = 1000, and ‘patience’ = 20. The encoder was subsequently frozen, and a linear classifier was trained on the sample representations with the following parameters: ‘lr’ = 3e-5, ‘weight_decay’ = 0.01, ‘max_epochs’ = 200, and ‘patience’ = 20. Sample representations were obtained for held-out samples using the trained model.

#### scGPT

scGPT was implemented following the authors’ official documentation (https://scgpt.readthedocs.io/). Cell embeddings were extracted from the pretrained scGPT model using the scg.tasks.embed_data function with the following parameters: ‘gene_col’ = “index”, ‘batch_size’ = 256, and ‘use_fast_transformer’ = True. The cell embeddings within each cell type were mean pooled, and the resulting cell type embeddings were averaged to obtain the sample representation. A linear classifier was trained on the sample representations with the following parameters: ‘lr’ = 3e-5, ‘weight_decay’ = 0.01, ‘max_epochs’ = 200, and ‘patience’ = 20.

#### tGPT

tGPT was implemented following the authors’ official documentation (https://github.com/deeplearningplus/tGPT). Raw count matrices were converted to ranked gene sequences using the process_adata_to_sequences function. The pretrained tGPT model was loaded using the GPT2LMHeadModel.from_pretrained. The tokenizer was loaded using PreTrainedTokenizerFast.from_pretrained function, and the sequences were tokenized with the following parameters: ‘max_length’ = 128, ‘truncation’ = True, and ‘padding’ = True. Cell embeddings were extracted by mean pooling the last hidden states from token position 1 to the end of the sequence. Cell embeddings within each cell type were averaged. The cell type representations were then averaged to obtain the sample representation. A linear classifier was trained on the sample representations with the following parameters: ‘lr’ = 3e-5, ‘weight_decay’ = 0.01, ‘max_epochs’ = 200, and ‘patience’ = 20.

#### scVI

scVI was implemented following the authors’ official documentation (https://docs.scvi-tools.org/). A total of 1200 highly variable genes were selected from the data using the Seurat v3 method. The data object was set up using the scvi.model.SCVI.setup_anndata. The model was initialized using the scvi.model.SCVI function with ‘n_latent’ = 32, and trained with scvi_model.train function. The cell embeddings were extracted using the get_latent_representation function. Cell embeddings within each cell type were averaged, and the cell type representations were then averaged across cell types to obtain the sample representation. A linear classifier was trained on the sample representations with the following parameters: ‘lr’ = 3e-5, ‘weight_decay’ = 0.01, ‘max_epochs’ = 200, and ‘patience’ = 20.

#### Simple multi-layer perceptron

Simple multi-layer perceptron (Simple MLP) model was implemented as a baseline for extracting sample representations. Raw count matrix was log1p-transformed and z-score scaled. The model was constructed with a two-layer multilayer perceptron architecture with GELU activation and a linear classifier. The model was trained with cross-entropy classification loss with the following parameters: ‘embedding_dim’ = 128, ‘hidden_dim’ = 512, ‘lr’ = 1e-5, ‘weight_decay’ = 0.01, ‘max_epochs’ = 400, and ‘patience’ = 10. Cell embeddings within each cell type were averaged, and the cell type representations were then averaged across cell types to obtain the sample representation. Sample representations were obtained for held-out samples using the trained model.

#### Principal component analysis

Principal component analysis (PCA) was implemented as a baseline for extracting sample representations. A total of 2000 highly variable genes were selected from the training data using the Seurat v3 method. Raw count matrices were log1p-transformed and z-score scaled. Principal components were extracted using the sklearn.decomposition.PCA function with ‘n_components’ = 32. Cell embeddings within each cell type were averaged, and the cell type representations were then averaged to obtain the sample representation. A linear classifier was trained on the sample representations with the following parameters: ‘lr’ = 3e-5, ‘weight_decay’ = 0.01, ‘max_epochs’ = 200, and ‘patience’ = 20.

### Evaluation

#### Sample-level disease prediction

We used five-time repeated random subsampling validation, where the data were randomly partitioned into train and held-out sets at the sample level, stratified by phenotype label. Each method was trained on the training set and evaluated on held-out samples. We then computed performance across the five iterations using the following metrics:

##### AU-ROC

The area under the receiver operating characteristic curve (AU-ROC) measures the ability of the classifier to discriminate between sample phenotype classes across all classification thresholds. It was computed as:

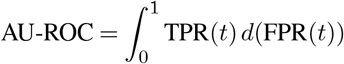

where TPR(*t*) and FPR(*t*) are the true positive rate and false positive rate at threshold *t*, respectively.

##### AU-PRC

The area under the precision-recall curve (AU-PRC) quantifies the balance between precision and recall across all classification thresholds. We used a one-versus-rest approach to obtain AU-PRC separately for each phenotype class. It was computed as:

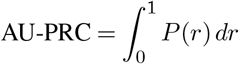

where *P* (*r*) is the precision at recall *r*.

#### Inference of disease trajectories

We evaluated whether the inferred trajectories recover the clinical and pathological variations using diffusion pseudotime (DPT) (63, 64). A k-nearest neighbor graph was built in the sample representation space, and a diffusion map was obtained. Subsequently, pseudotime was computed from the resulting diffusion components. For the COVID-19 dataset (21), a control sample was randomly selected as the trajectory root, and severity categories (control, mild/moderate, and severe/critical) were encoded as ordinal values. The Spearman correlation was then computed between inferred pseudotime and encoded severity. We additionally fit an ordinal logistic regression model with pseudotime as the predictor and disease severity as the ordered outcome. In the Alzheimer’s disease dataset (26), the reference sample with the lowest neuropathological burden was set as the trajectory root. We then computed Spearman and Pearson correlations between inferred pseudotime and pathological measures.

#### Alzheimer’s disease analysis

We used the Phenoverse inferred trajectories to characterize cross-cohort molecular programs in the ROSMAP and SEA-AD Alzheimer’s disease cohorts. In the ROSMAP cohort, the sample with the lowest neuropathological burden was chosen as the trajectory root, and the disease trajectory was inferred. In the SEA-AD cohort, a sample annotated as reference was randomly selected as the root to infer the trajectory. We computed per-cell type pseudobulk gene expression profiles per sample by summing raw counts. Within each cell type, genes were retained if expressed in ≥ 1% of cells and with CPM *>* 0.5 in ≥ 30% of samples.

For trajectory genes, differential expression was conducted using pyDESEQ2 with pseudotime as a continuous design factor (32, 33). For the binary case-control comparison, sample disease labels in each cohort were used to perform differential expression analysis with pyDESEQ2. We quantified the cross-cohort reproducibility of different gene rankings for each cell type by computing the Spearman correlation of gene-level Wald statistics across shared genes. The pathway activity along the Phenoverse inferred trajectory was computed per cell type by performing single-sample gene set enrichment analysis (ssGSEA) against hallmark gene sets from the molecular signatures database (MSigDB) (34–37). The Spearman correlation between inferred trajectory and pathway activity scores was computed across samples. The p-values were then corrected using the Benjamini-Hochberg false discovery rate (FDR) approach. Pathways with Spearman |*ρ*| ≥ 0.25 and FDR *<* 0.05 across both cohorts with concordant directionality were classified as reproducibly enriched or depleted.

#### Systemic lupus erythematosus analysis

We used Phenoverse to identify cell type-specific disease states in the SLE single-cell transcriptomic cohort (22). A k-nearest neighbor graph was constructed from sample representations, and Leiden community detection was applied (resolution = 0.5) to identify clusters (65). To quantify interferon activity per sample, cells were log-normalized to 104 counts per cell, and a per-cell IFN score was computed using a panel of 16 interferon-stimulated genes (ISGs) with scanpy (64). We then computed the per-sample IFN score by averaging across all cells for each sample. For each cell type, a Kruskal-Wallis test was performed per prototype to assess the occupancy across clusters. The p-values obtained were corrected using the Benjamini-Hochberg FDR approach.

The prototype direction was determined by computing the Spearman correlation between the cluster mean prototype occupancy and the cluster IFN rank. Prototypes with Kruskal-Wallis FDR *<* 0.05 and direction Spearman |*ρ*| ≥ 0.3 were classified as disease-associated prototypes. For each cell type, genes expressed in ≥ 1% of cells were retained. A Spearman correlation was computed between per-cell log-normalized gene expression and per-cell prototype occupancy. The p-values were corrected using the Benjamini-Hochberg FDR approach. For each cell type, genes expressed in ≥ 1% of cells were used as the background, and gene set enrichment analysis was performed against MSigDB Hallmark gene sets for the top 200 positively correlated genes (FDR *<* 0.05) per prototype (36, 37, 66).

## Data availability

The Phenoverse framework is implemented as a Python PyPI package, and the source code is available at https://github.com/KellisLab/Phenoverse. Documentation for using the tool is available at https://KellisLab.github.io/Phenoverse/.

## Acknowledgments

The authors thank the MIT Office of Research Computing and Data and the Broad Institute of MIT and Harvard for providing the high-performance computing resources that have contributed to the research results reported within this paper. M.M.W. is supported by scholarships from the University of Sydney and the Children’s Medical Research Institute. M.K. is supported by the Cure Alzheimer’s Fund and the Biswas Family Foundation Transformative Computational Biology Grant. P.Y. is supported by an Australian Research Council (ARC) Future Fellowship (FT250100252). We thank Amy Grayson, Li-Lun Ho, and Nicola Sykes for their support throughout the project. The authors also thank the intellectual engagement and feedback from colleagues at the MIT Computer Science & Artificial Intelligence Laboratory, the University of Sydney, and the Children’s Medical Research Institute. The authors used Anthropic’s Claude Sonnet 5 thinking model to assist with improving the writing during manuscript preparation. The authors reviewed and assume responsibility for the content of the article. The results published here are in whole or in part based on data obtained from the AD Knowledge Portal. ROSMAP data were generated from postmortem brain tissue provided by the Religious Orders Study and Rush Memory and Aging Project (ROSMAP) cohort at Rush Alzheimer’s Disease Center, Rush University Medical Center, Chicago. SEA-AD data were generated from postmortem brain tissue obtained from the University of Washington BioRepository and Integrated Neuropathology (BRaIN) laboratory and Precision Neuropathology Core, which is supported by the NIH grants for the UW Alzheimer’s Disease Research Center (P50AG005136 and P30AG066509) and the Adult Changes in Thought Study (U01AG006781 and U19AG066567).

## Author information

M.M.W. - Conceptualization, Data curation, Software, Formal analysis, Investigation, Methodology, Writing-original draft, Writing-review and editing; Y.W. - Conceptualization, Methodology; S.S. - Formal analysis; Z.L. - Data curation, Methodology; E.P. - Conceptualization, Supervision, Methodology, Project administration; P.Y. - Conceptualization, Supervision, Methodology, Project administration, Writing-review and editing; M.K. - Conceptualization, Resources, Supervision, Methodology, Project administration, Writing-review and editing.

---

**Supplementary Figure 1.**
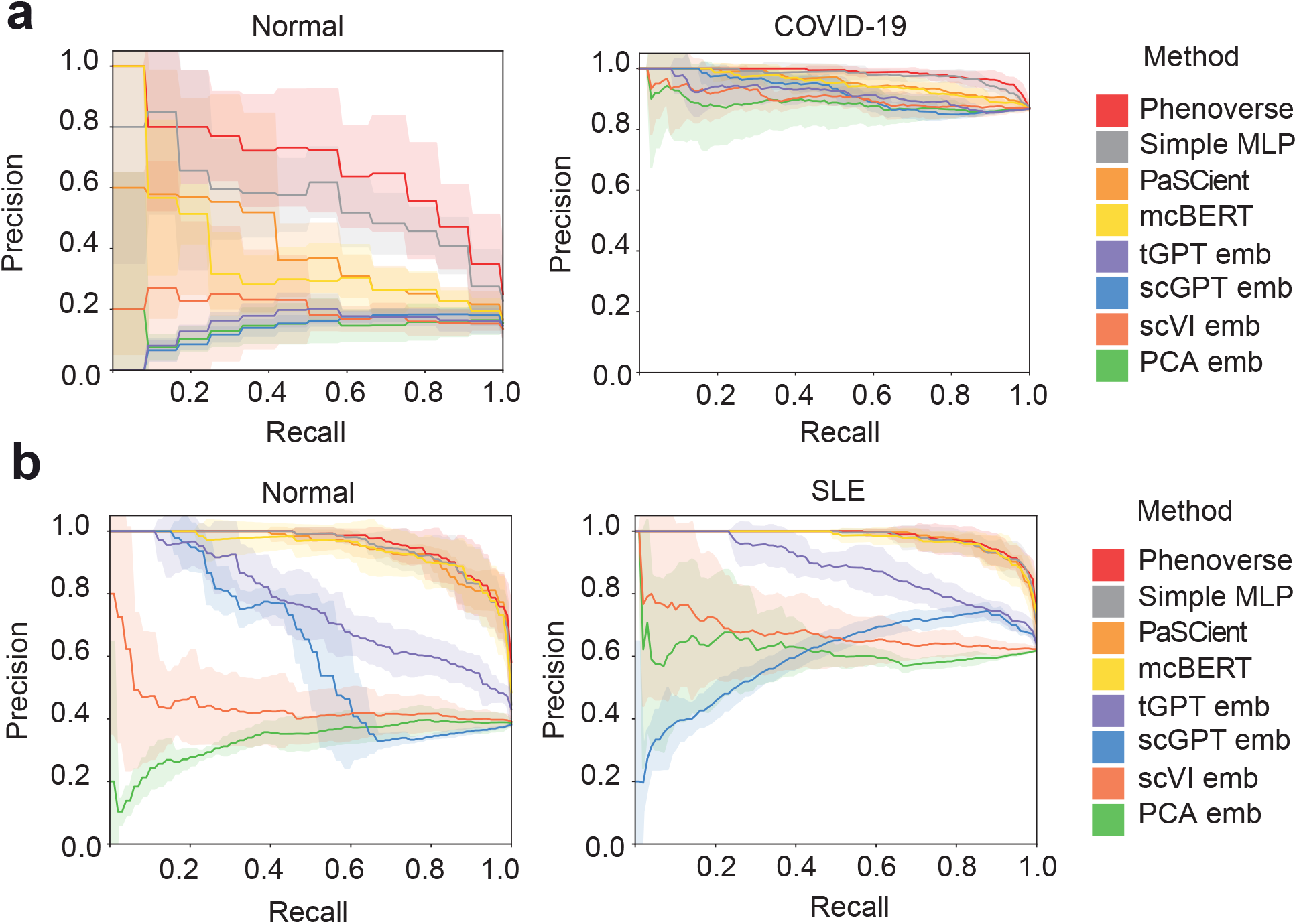
Disease state prediction performance of different methods. (a) Mean precision-recall curves across five repeated subsampling splits in the COVID-19 dataset, shown separately for the Normal (left) and COVID-19 (right) classes. Shaded bands indicate *±*1 standard deviation across splits. (b) Mean precision-recall curves across five repeated subsampling splits in the systemic lupus erythematosus (SLE) dataset, shown separately for the Normal (left) and SLE (right) classes. Shaded bands indicate *±*1 standard deviation across splits.

**Supplementary Figure 2.**
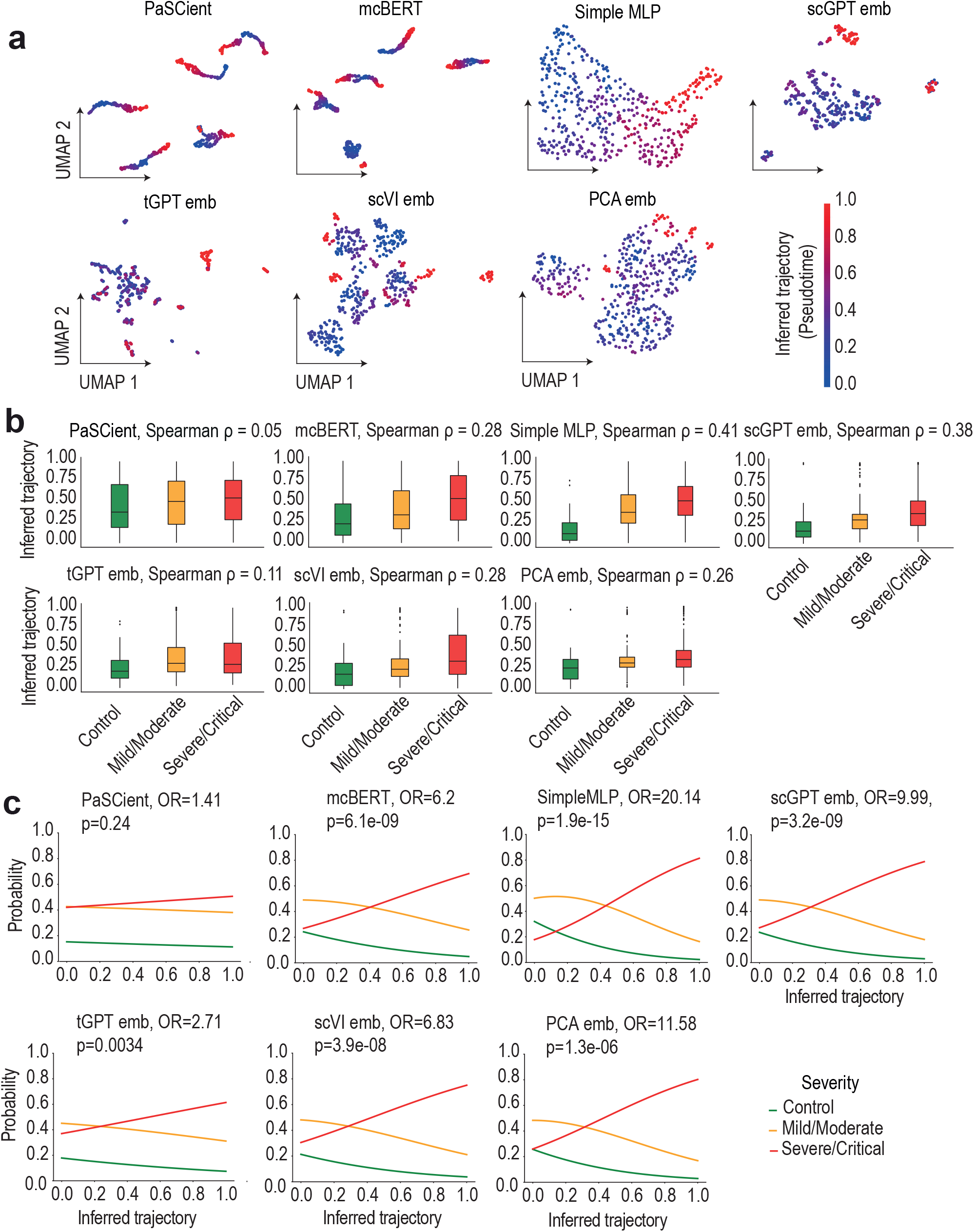
Disease trajectory inference performance of baseline methods in the COVID-19 dataset. (a) UMAP of sample representations for each baseline method from held-out test samples in the COVID-19 dataset (21), colored by inferred trajectory (pseudotime). (b) Box plots showing the distribution of inferred trajectories across clinical severity categories (control, mild/moderate, and severe/critical) for each baseline method. Box plots show the median (center line), interquartile range (IQR) box bounds: 25th–75th percentile, 1.5*×*IQR (whiskers), and points outside 1.5*×*IQR as outliers. (c) Line plots showing ordinal logistic regression predicted probabilities for each clinical severity category along the inferred trajectory for different baseline methods. Odds ratios (OR) and associated p-values are indicated in each panel title.

**Supplementary Figure 3.**
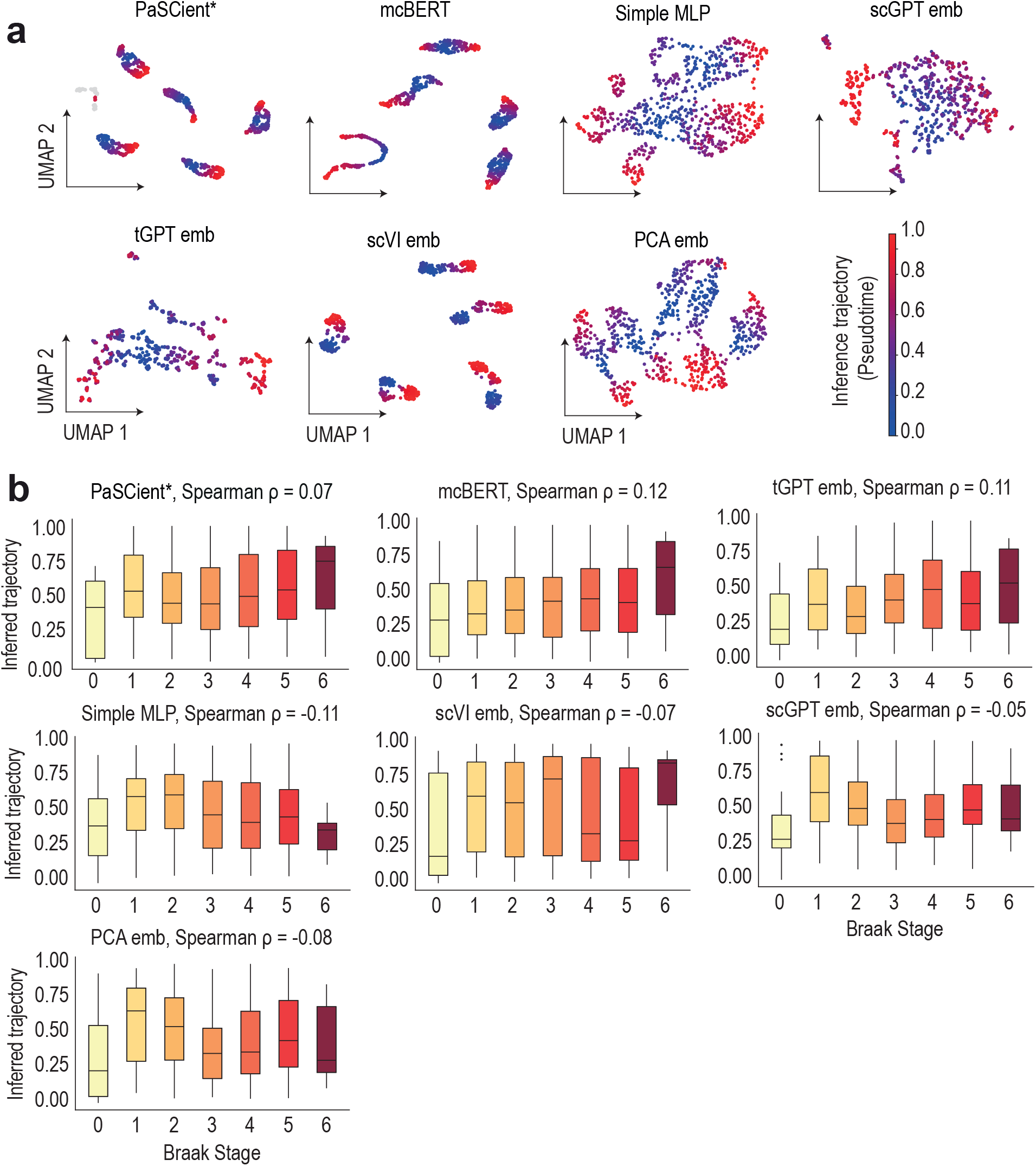
Correlation of inferred trajectories of baseline methods with Braak staging in Alzheimer’s disease. (a) UMAP of sample representations for each baseline method from held-out test samples (26), colored by inferred trajectory (pseudotime). * indicates that pseudotime values were undefined for a subset of held-out samples. (b) Box plots showing the distribution of inferred trajectories across Braak stages (0–6) for each baseline method. Box plots show the median (center line), interquartile range (IQR) box bounds: 25th–75th percentile, 1.5*×*IQR (whiskers), and points outside 1.5*×*IQR as outliers. * indicates that pseudotime values were undefined for a subset of held-out samples.

**Supplementary Figure 4.**
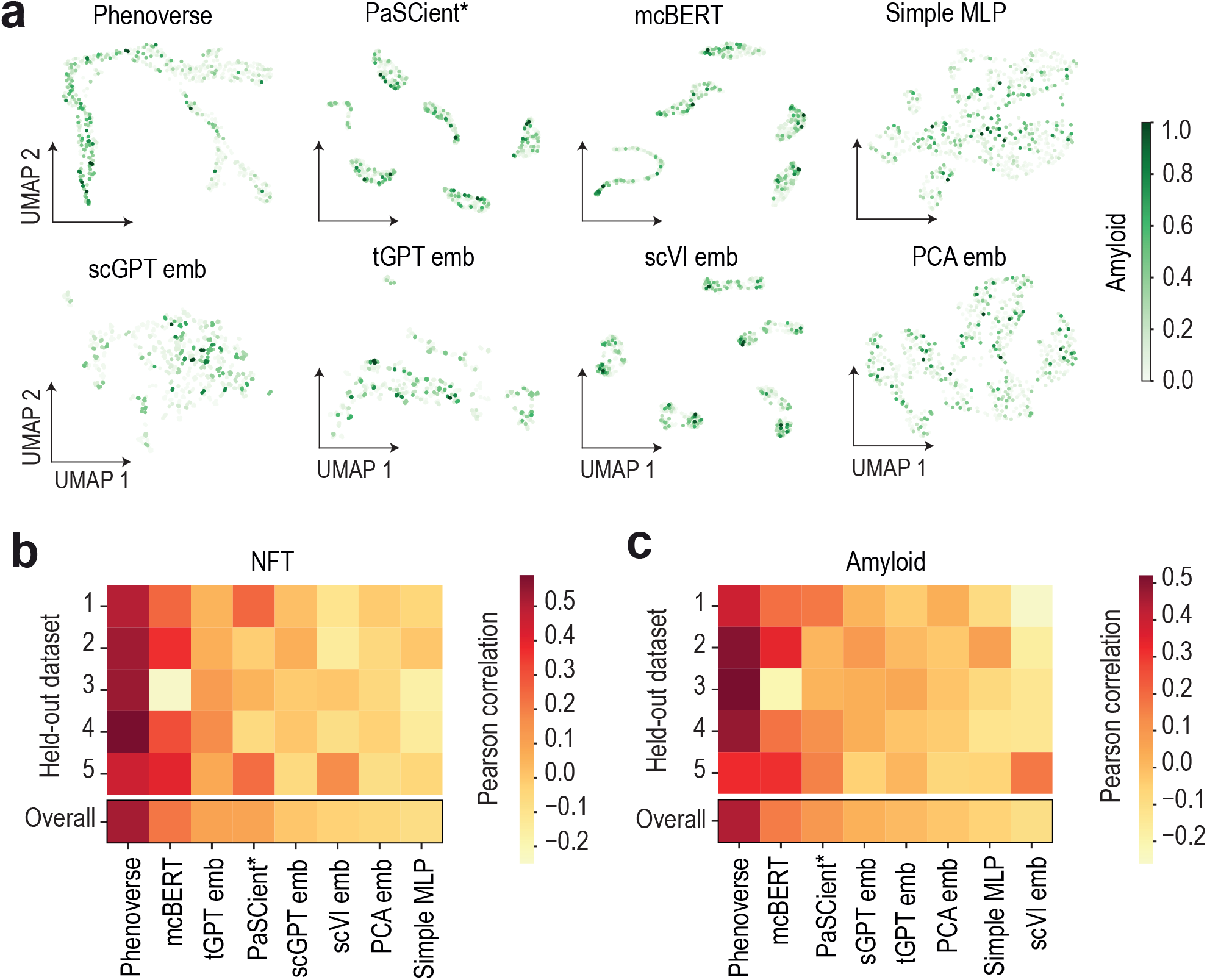
Correlation of inferred trajectories of baseline methods with neuropathological burden in Alzheimer’s disease. (a) UMAPs of sample representations from held-out test samples in the Alzheimer’s disease dataset (26), colored by amyloid burden. * indicates that pseudotime values were undefined for a subset of held-out samples. (b-c) Heatmaps showing the Pearson correlation between inferred trajectory and NFT burden (b) and amyloid burden (c) across five repeated subsampling validation splits. * indicates that pseudotime values were undefined for a subset of held-out samples.

**Supplementary Figure 5.**
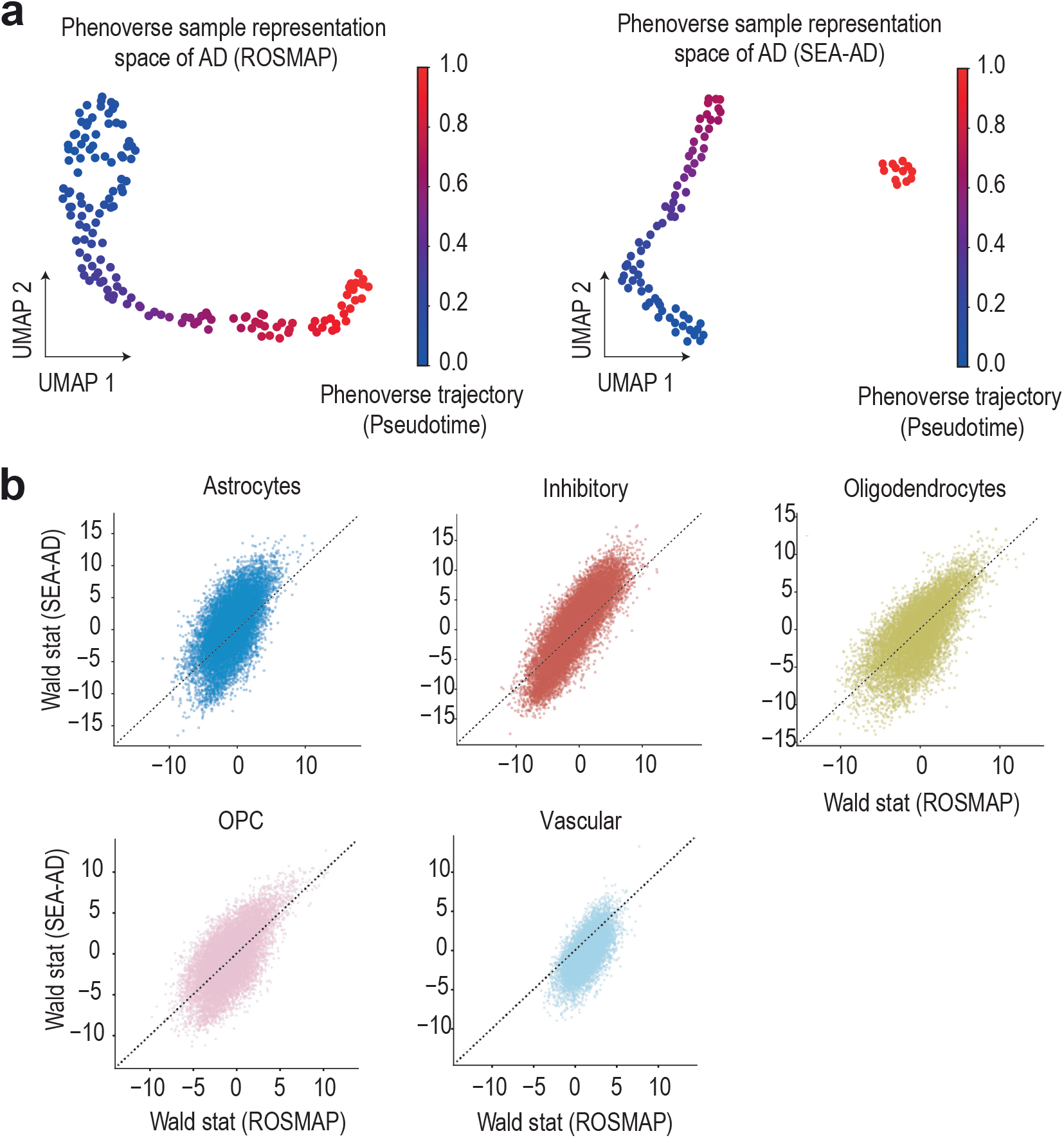
Cross-cohort trajectory inference and gene-level reproducibility in Alzheimer’s disease. (a) UMAPs of Phenoverse sample representations from held-out ROSMAP (left) and SEA-AD (right) samples, colored by inferred trajectory (pseudotime). (b) Scatter plots showing the cross-cohort reproducibility of gene-level Wald statistics from Phenoverse trajectory-based differential expression analysis across different cell types.

**Supplementary Figure 6.**
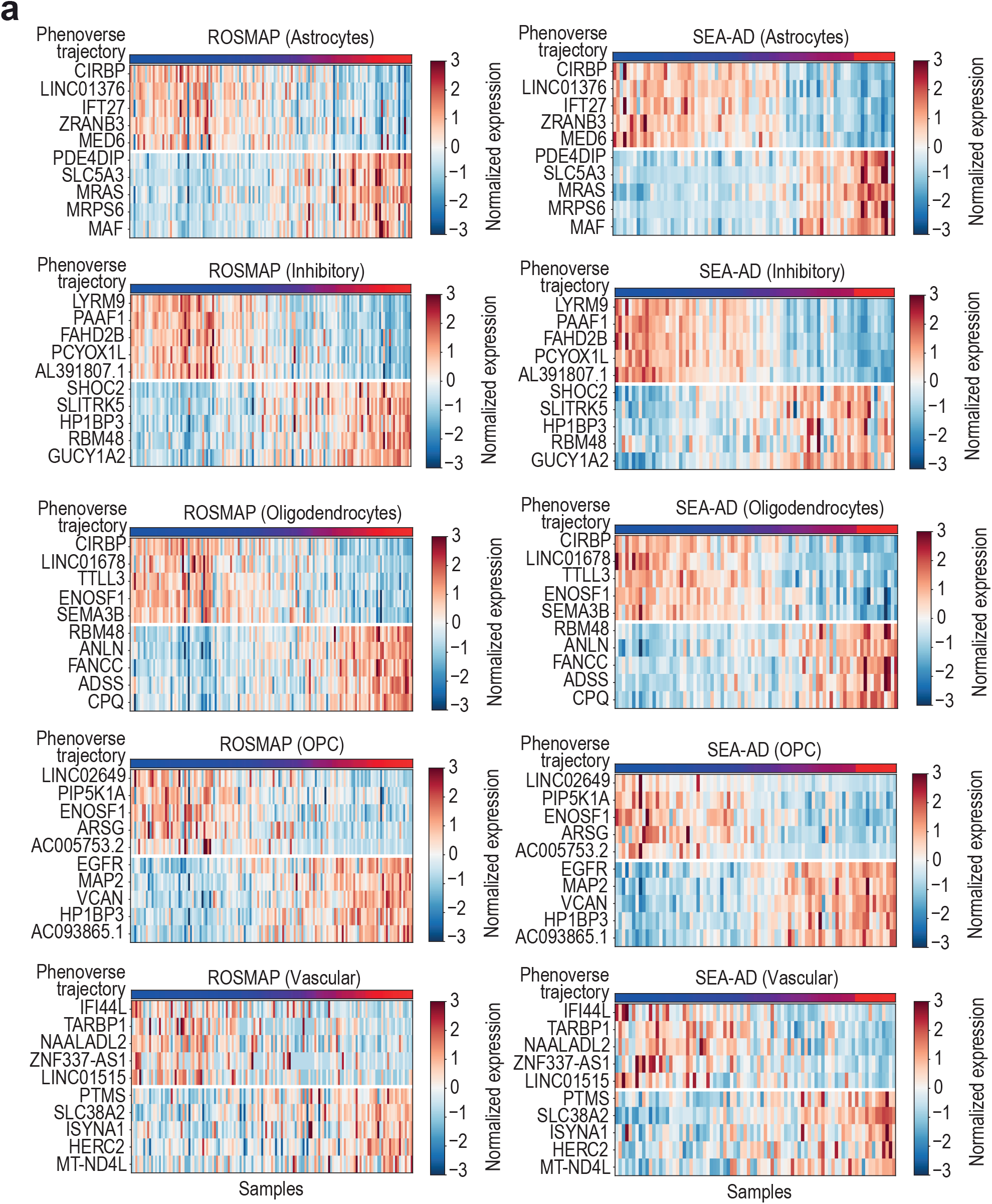
Expression patterns of Phenoverse trajectory-associated genes. (a) Heatmaps showing expression patterns of top upregulated and downregulated trajectory-associated genes across different cell types. Genes were selected based on Wald statistics from the ROSMAP cohort. Expression values are shown as row-normalized Z-scores for ROSMAP (left) and SEA-AD (right) samples. Rows correspond to genes, and columns correspond to samples ordered by increasing pseudotime.

**Supplementary Figure 7.**
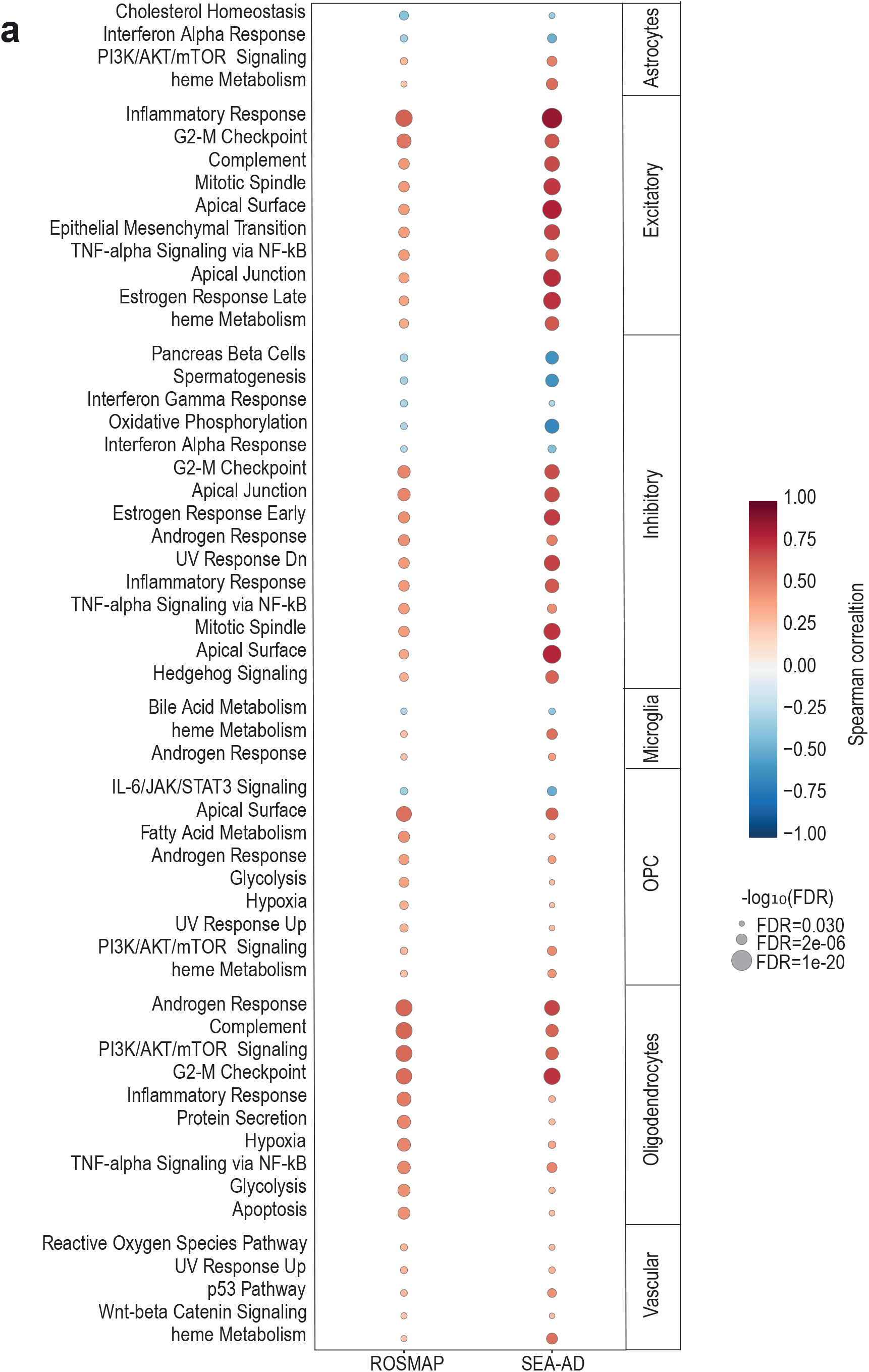
Molecular programs along Phenoverse-inferred trajectories in Alzheimer’s disease. (a) Dot plot showing the top 10 significant hallmark pathways along Phenoverse-inferred trajectories across seven cell types in ROSMAP and SEA-AD cohorts. Dot size represents the magnitude of the Spearman correlation between pathway activity scores and the Phenoverse-inferred trajectory. Color indicates direction relative to the inferred trajectory (red: enriched; blue: depleted).

**Supplementary Figure 8.**
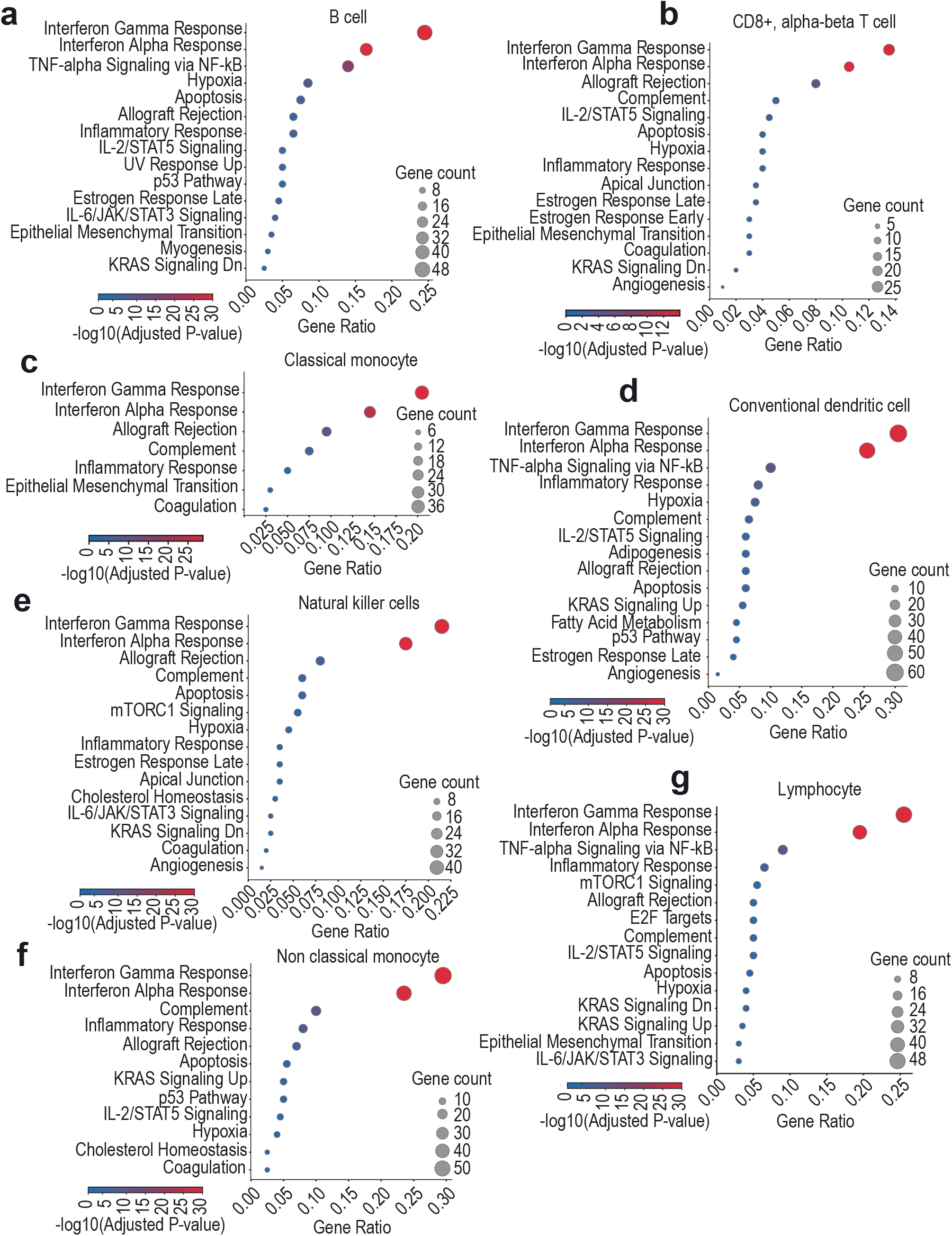
Prototype enriched programs in systemic lupus erythematosus. (a-g) Dot plots illustrating enrichment of top prototype 3 gene programs in B cell (a), CD8+ alpha-beta T cell (b), classical monocyte (c), conventional dendritic cell (d), natural killer cell (e), non-classical monocyte (f), and lymphocyte (g).

